# Design, construction, and advantages of 100% online laboratories in an upper division undergraduate biology course

**DOI:** 10.1101/2020.12.30.424788

**Authors:** Emma Brady, Shalaunda Reeves, Malcolm Maden, Brian D. Harfe

## Abstract

For centuries, students have been taught by passively listening to lectures, usually in large groups. In many fields, this type of course delivery is not ideal. In addition, students that have grown up with online access and/or have been forced to take online course during the recent pandemic, are seeking improved online educational experiences. As providers of education, Universities must adapt to this rapidly changing environment. The emergence of exciting, “disruptive” technologies has provided the tools needed for educators to revolutionize the delivery of higher education. The University of Florida, through UF Online, provides students the opportunity to enroll in fully online four-year undergraduate degrees. In this report, we describe the development of fully online laboratories for the upper-division course Evolutionary Developmental Biology, which fulfill the University of Florida’s laboratory requirement. This course is taken by Biology, Zoology, and related majors, including majors in the 100% online Biology degree. The experiments in the virtual laboratories would have been impossible to teach in a residential setting due to time constraints, cost, student safety issues, and regulatory/certification requirements. The ability to perform these experiments online provided University of Florida undergraduates located throughout the world access to unique, one-of-a-kind laboratory experiences. The virtual laboratories allowed students to undertake in-depth exploration and application of course content that is not possible in a residential setting. Completion of the Evolutionary Developmental Biology course provided undergraduate students the opportunity to obtain a science degree without ever visiting the university’s residential campus. During the Fall 2020 term, this course was made available to residential students, thus allowing this cohort of students to continue to make progress, 100% online, towards their undergraduate degree during the COVID19 pandemic.

## Introduction

Laboratories are a conventional component of undergraduate degree programs, especially in biology courses that fulfill a general education requirement. While the development and teaching of online lower division biology laboratories has gained momentum, the development of upper division biology laboratories for biology and related majors has proven problematic [1, 2]. This is likely due to a number of factors, including the belief that it is inappropriate to provide online laboratories to science majors, the technological difficulty in designing realistic online laboratory experiences, and the potential cost and time it takes to develop online laboratories[1]. In this report, we will describe our experience developing fully online laboratories, address some of the key issues in the field of online laboratory development, and discuss the advantage of having robust online offerings during a pandemic.

In the fall of 2013, in response to the Florida Senate Appropriations subcommittee on Education[3], the University of Florida (UF) was directed by the Florida Legislature to create fully online undergraduate degrees to address issues related to student accessibility in high education. By Florida state statue, these “UF Online” students are prohibited from taking residential classes on the University of Florida campus. Students pay 75% of the tuition cost of residential students and three of the seven residentially required fees to attend the University of Florida (unless online students choose optional additional services). UF Online students are required to meet the same admission and degree requirements as residential students. Upon completion of their degree, UF Online students receive diplomas and transcripts that are identical to University of Florida residential students.

Upon creation of the online degree program by the Florida legislature, an exemption to the “100% online” requirement was granted for laboratory courses. This exemption provided the possibility for UF Online students to find an equivalent residential laboratory at a local community college or university. However, providing only an in-person option for UF Online students creates a significant barrier to graduation for a student body that depends on the flexibility of an online degree. To address this barrier, the Biology Department in the UF College of Liberal Arts and Sciences and UF Online decided to develop an online version of the upper division course Evolutionary Development Biology (course number ZOO3603C) that contained three fully embedded online laboratories. This allowed the entire course to be taken virtually.

## Description of the course Evolutionary Developmental Biology (ZOO3603C)

Evolutionary Developmental Biology is an upper division elective course for Biology and related majors. Students must have completed the two-semester biology introductory course sequence for Biology majors, prior to enrolling in Evolutionary Developmental Biology. Both the residential and 100% online versions of this course are taught every fall and each usually enrolls ~40 junior and senior Biology/Microbiology/Zoology majors. Residential and online versions of the course include laboratories as well as lectures.

Evolutionary Developmental Biology is designed to provide students an understanding of how an organism develops by studying the developmental principles observed in various organisms. Model systems described in the course include, invertebrates such as the fruit fly, lower vertebrates such as fish and frogs, and higher vertebrates such as birds and mammals. The development of individual organ systems such as the brain, eye, and the limbs are used to draw together principles of body organization. By studying the development of these different animal systems, students draw together principles of development which have stood the test of evolutionary time. In addition, the course also includes a consideration of the regeneration of complex organ systems such as the limb and the molecular pathways involved in these processes.

## Virtual laboratory development and integration

This report will focus on the development and implementation of three first-person, virtual laboratory simulations, developed in collaboration with Labster, a virtual laboratory development company located in Copenhagen, Denmark. The result of this unique collaboration was the creation of laboratories that fulfill the University of Florida’s laboratory requirement for an upper division course for Biology and related majors. This course is one of the first online upper division Biology major course that includes 100% online laboratories taught at an Association of American University (AAU).

The laboratories were strategically embedded into the online lecture version of ZOO3603C, which was developed simultaneously. Lectures in the course were recorded using a “green screen”. The course piloted in the fall 2016 semester and is currently offered each fall. The laboratories were designed to encourage critical thinking related to applying course content and to provide practice in standard laboratory procedures. Each of the laboratories were designed to demonstrate and explore the unique advantages of different animal model systems. The first laboratory, “Discovering genes responsible for patterning a vertebrate limb” used the mouse and chick model systems; the second laboratory, “The power of *C. elegans* in human gene discovery”, used the *C. elegans* model system; and the third laboratory, “Investigation of regenerative capabilities” explored the axolotl model system.

Laboratories were completed in weeks 4, 11, and 15 of the typical 16-week semester. During the weeks prior to each laboratory, online lectures, assignments, and discussions introduced and developed supporting course concepts including the advantages and disadvantages of each model system. During laboratory weeks, the course instructor presented a short introductory video designed to focus students’ attention on content application and what they were expected to accomplish in the laboratory. During the weeks that the laboratories were assigned, no new lecture material was presented.

## Construction of the vertebrate embryology laboratory using the mouse and chick model systems

Unlike many virtual laboratory simulations currently available on the market, the content for the three laboratories was developed entirely by active research faculty specifically to fulfill the objectives of the ZOO3603C course. Two of the authors of this report, (M.M and B.D.H) are Principle Investigators of developmental biology research laboratories and have published extensively in this field. They worked closely with developers at Labster to ensure that the simulations represented, as closely as possible, what occurs in a live research setting. For example, the close collaboration ensured that everything from equipment placement to the order of experiments was accurately portrayed in the virtual laboratory space. The ability of the team to create virtual laboratory simulations mitigate many of the limitations associated with performing similar experiments in a live residential setting. The initial development step for each laboratory was the creation of learning objectives. In the Vertebrate Embryology Laboratory, the three learning objectives were:

1. Compare and contrast the advantages and disadvantages of the mouse and chicken model systems.
2. Uncover a molecular pathway responsible for forming a vertebrate fore- or hindlimb.
3. Formulate and test a hypothesis investigating genetic causes of limb defects.

The vertebrate embryology laboratory begins in a hospital room where a patient with Liebenberg Syndrome is being examined by the student and a medical doctor. Liebenberg Syndrome is a unique condition where the arms of a person resemble their legs[4]. The transmission of the disease is autosomal dominant and mutations that can cause this disease has recently been identified[5–7]. However, students at this point in the laboratory were not informed that the underlying cause of this disease was known. The doctor and student take a case history of the patient where it becomes clear that multiple people in the patient’s family are affected. The introduction to this laboratory ends with the patient requesting that the student, along with their mentor, return to their laboratory to attempt to understand the genetic basis of the patient’s condition.

In the virtual laboratory, the student is asked by the doctor a series of multiple choice questions that are designed to demonstrate that a vertebrate model system is the best choice to investigate vertebrate limb development. The student is then directed to “window” a chicken egg (i.e. create an opening on the top of a fertilized chicken egg) using a dissecting microscope. The windowed egg allowed the student to examine a living embryo (Fig. 1a, b). By placing their egg in an incubator and removing it at various time points the student was able to watch their egg undergo development.

**Figure 1.**
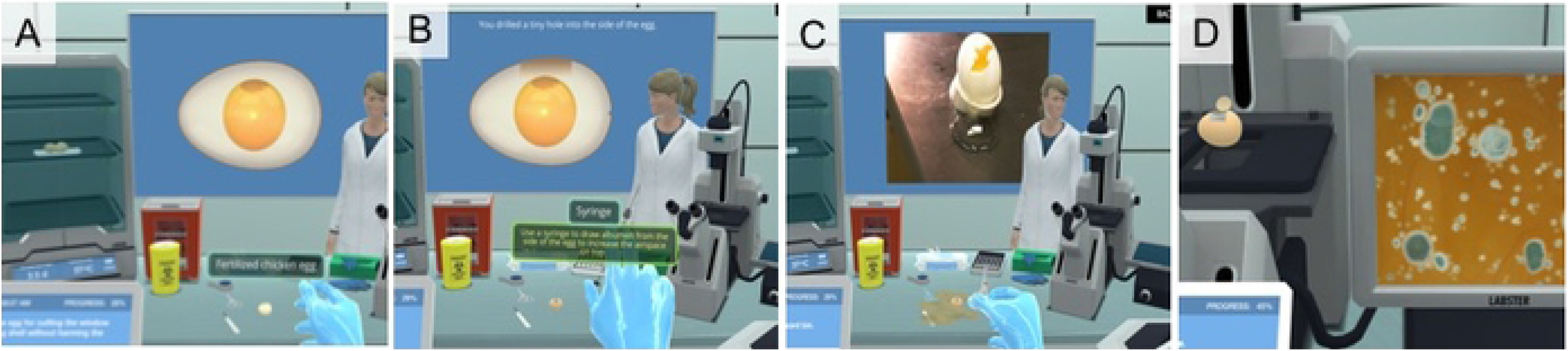
Images taken from Laboratory 1, “Discovering genes responsible for patterning a vertebrate limb”. (A) By placing eggs in an incubator and removing them at various time points students were able to watch their eggs undergo embryonic development. (B) The student decides when and where to use a syringe to lower albumen levels prior to removing the egg shell during “windowing” of the egg. (C) Incorrect windowing resulted in albumen leaking out of the egg. If the egg was contaminated, as seen in (D), the student was able to immediately repeat the windowing experiment with no loss in time or resources.

These experiments demonstrate two key advantages of performing experiments in an online environment. The first is the ability to speed up (or slow down) time. The analysis of different stages of embryonic development would normally require a student to place their egg in an incubator and then return to the laboratory every ~2-3 days to examine development. Since most laboratories only meet once a week and/or do not have the resources or time to perform multi-week experiments, it can be difficult to examine development of a vertebrate organism over an extended time-period in a residential laboratory setting. While this limitation can be overcome by a laboratory manager preparing a “staged” series of embryos at different time points prior to students arriving at the laboratory, this does not allow students to examine their individually windowed egg.

Online laboratories have been criticized for their inability to allow students to make “mistakes”[8]. While mistakes and accidents are a key component of learning, these can also cause a laboratory experiment to be halted prior to all learning outcomes associated with a given laboratory having been completed. The egg windowing experiment is an excellent example of how an online laboratory can allow students to “mess-up” an experiment while maintaining the student’s ability to proceed through all the learning objectives. For example, the online windowing experiment has two potential areas where a student may make a mistake. The first is the placement of the needle in the egg, which is required to lower albumen levels prior to removing the eggshell. Students that incorrectly perform this procedure cause albumen and the embryo to leak out of the egg, identical to what occurs in a residential setting (Figure 1c). In both online and residential laboratories, this results in loss of the embryo and the need to repeat the experiment. However, embryo loss in the online environment is inconsequential, in regards to resources and time, allowing the student to immediately repeat the experiment.

A second common issue with performing experiments using chicken eggs is bacterial and/or fungal contamination of the egg leading to death of the embryo. Bacterial contamination commonly occurs if a student does not sterilize the outside of the egg prior to windowing. However, death of the embryo does not occur immediately. Only after examination of the embryo after the desired time points (usually in ~48 hour increments) would a student realize that their egg had become contaminated. In the residential setting, death of the embryo results in the halting of the experiment (unless multiple eggs were used by each student and at least one survived to the required embryonic stage. The use of multiple eggs per student increases the cost of the laboratory). In the online setting, students do not have to wait to see the results of their experiment. Students put their windowed eggs in the incubator, speed up time, and then examine the results. If their egg was contaminated (Fig. 1d), they can immediately repeat the windowing experiment with no loss in time or resources. Alleviating the limitations of cost and time allowed student to repeat experiments as often as they needed to obtain mastery of the material.

To determine the stage of the embryo being examined, students viewed a three-dimensional model of a chicken embryo. By sliding a progress bar beneath the model, the student could increase or decrease the age of the embryo. Once they identified a stage that matched the stage of their windowed egg, they could determine the exact age of their embryo. During this part of the laboratory, students were prompted to pay particular attention to the limbs, since their goal was to find an embryonic stage where genes involved in Liebenberg Syndrome may play a role in the patterning of this structure.

The second part of the laboratory used the mouse model system to uncover genes that were potentially involved in causing Liebenberg Syndrome. The mouse model was introduced by examining a mouse embryo under the virtual microscope and asking the student if the embryo was a chicken or mouse embryo. The student was then asked what human embryonic stage would match the stage of mouse limb development shown. This example of inter-species comparison enforced the concept that model systems are ideal for investigating human diseases since chick, mouse, and human limb development closely resemble each other, a concept integral to the overall course goals.

To determine genes that, upon mutation, were potentially involved in arms resembling legs in humans (a characteristic of Liebenberg Syndrome[4]), mouse fore- and hindlimbs were dissected from embryos. The hypothesis being tested was that a gene(s) required for the development of these structures may be enriched in the forelimb or hindlimb. Genes expressed equally in both regions would not be candidates for the specification of the unique characteristics that distinguished fore- and hindlimbs.

The dissection of mouse limbs is technically challenging and time consuming. In the published literature, including a paper from one of the authors of this report[9], harvesting of mouse limbs for differential gene analysis requires multiple litters of animals. In addition, training is required for a student to be able to consistently dissect limb tissue from the flank of a, relatively, small embryo under a dissecting microscope. Students that perform these experiments in a residential setting would be required to undertake Institutional Animal Care and Use Committee (IACUC) training prior to working with vertebrate animals (Table 1). Due to these constraints, the use of the mouse model system in an undergraduate laboratory setting is extremely rare.

**Table 1.** Training that would be required at the University of Florida to perform the experiments present in the three online laboratories if they were provided in a brick-and-mortar environment.

| Technique | Training |
| --- | --- |
| Performing experiments with mice | IACUC – Working with Vertebrate Animals |
| Performing experiments with axolotls | IACUC – Working with Vertebrate Animals |

To determine gene expression differences in the fore- and hindlimbs, Next Generation Sequencing was performed in the online laboratory. This technique generates short sequencing “reads” of billions of RNA fragments generated from transcribed genes that were present in the limb buds when they were harvested. Genes that were, for example, expressed at higher levels in the hindlimb relative to the forelimb can be identified using this technique. To determine genes that were differentially expressed, the student was promoted to overlay Next Generation Sequencing data showing the number of times a given gene transcript was present in either the fore- or hindlimbs. These data uncovered two genes, *Pitx1* and *Tbx4*, that were enriched in the hindlimb compared to the forelimb, making them excellent candidates for causing Liebenberg Syndrome. Both, *Pitx1* and *Tbx4*, have been demonstrated to play essential roles in limb development and patterning[10–12].

To conclude the laboratory, the student sequences human homologs of *Pitx1* and *Tbx4* from the patient they took a family history from at the beginning of the laboratory. The student is told that the DNA sequencing data revealed the presence of a mutation near *Pitx1* in the human patient and that the identified mutation is in a region of the gene that may cause the gene to be aberrantly expressed in the forelimb (it is normally only expressed in the hindlimb). In humans, mutations in *Pitx1* are associated with Liebenberg Syndrome[5–7, 13]. These data suggest that these mutations may be responsible for Liebenberg Syndrome. Finally, the student is asked what further experiments could be performed to test the hypothesis that the gene they identified may be causing arms to resemble legs.

## Construction of the invertebrate genetics laboratory using the C. elegans model system

In the invertebrate embryology laboratory, the learning objectives were:

1. Determine characteristics of invertebrate model systems that make these animals ideal for uncovering genes involved in human diseases.
2. Distinguish between male and hermaphrodite *C. elegans*.
3. Identify and characterize mutant *C. elegans* generated in a forward genetic screen.

The laboratory begins with the student observing a case presentation in a hospital conference room. A three-year-old child is shown who has dysmorphic facial features consistent with Saethre Chotzen Syndrome. Saethre Chotzen Syndrome is an autosomal dominant disease (one mutated copy of a gene results in manifestation of the disease)[14]. Approximately half of all cases of Saethre Chotzen Syndrome are due to mutation of a gene called *twist[15, 16]*. TWIST is a transcription factor that can turn on other genes during embryonic development. The child presented in the laboratory does not have a *twist* mutation, indicating that other genes, upon mutation, can cause Saethre Chotzen Syndrome. The scene then shifts to the laboratory, where the student begins their experiments to identify what mutations, besides mutations in *twist*, could cause Saethre Chotzen Syndrome.

The model organism used in this laboratory are *C. elegans*. *C. elegans* are small, ~1mm, nematodes usually found in soil, that have been used extensively in laboratories throughout the world to identify gene function [17]. The student, through correctly answering multiple choice questions in the online laboratory, are taught some of the key characteristics of *C. elegans*. In addition, prior to beginning the laboratory, students complete the *C. elegans* lecture module of the course. This module provided background on many of the key characteristics[18] that have made *C. elegans* popular as a laboratory model system.

*C. elegans* have two sexes, hermaphrodites and males. To perform genetic crosses using *C. elegans* it is essential that a student be able to visually identify the two sexes as well as the different larval stages of development. In the first experiment, students visualize *C. elegans* development using a dissecting microscope and then “pick” animals by clicking on them (Figure 2a). After successful identification of all stages of *C. elegans* development, the student begins the genetic screen.

**Figure 2.**
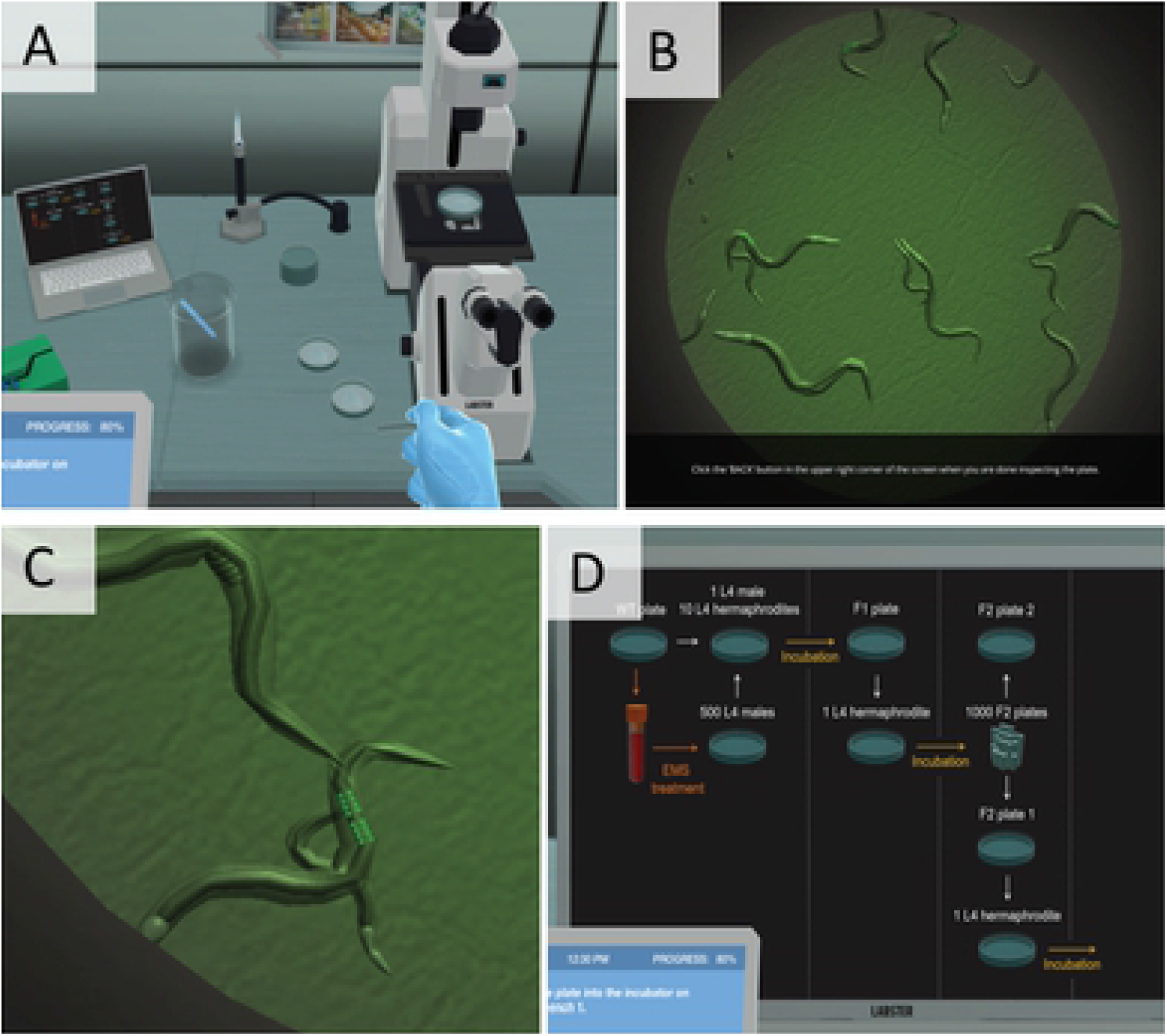
Images taken from Laboratory 2, “The power of *C. elegans* in human gene discovery”. (A) Students visualized *C. elegans* development using a dissecting microscope. (B) Animals were “picked” by clicking on them. (C) Zoomed picture of C. elegans in which the GFP-positive M cells can be seen (white arrow). (D) To aid students in understanding the genetic crosses they needed to perform, a real-time flow chart was filled in on a virtual laptop as student’s progress through the laboratory.

*C. elegans* are transparent, which allows researchers to easily visualize a florescent protein call Green Florescent Protein (GFP) in individual *C. elegans* cells (one third of the 2006 Nobel Prize was awarded to Dr. Martin Chalfie for his use of GFP in *C. elegans*)[19]. In the virtual laboratory, students are provided *C. elegans* that express GFP in all cells of a specific muscle lineage called “M”. These GFP-positive M lineage cells also express *twist[20]*, one of the genes know to result in Saethre Chotzen Syndrome if mutated in humans *[15, 16].* Using a florescent dissecting microscope, which would typically cost approximately >$8,000 (Table 2), the student can observe these cells since they glow green due to the presence of GFP protein (Figure 2b, c). Students expose *twist*:*gfp C. elegans* to a mutagen and then identify animals that express GFP in an altered pattern. Animals with altered GFP expression may contain a mutation in a gene that works in the same pathway and/or patterns the same tissue as *twist*.

**Table 2.** Cost of major equipment used in the three laboratories. The costs below are for each individual laboratory and do not take into consideration savings that could be realized by reusing equipment common between laboratories. Costs are the approximate “list price” of equipment and does not represent potential educational or bulk discounts that may be provided by venders. The cost of Next Generation Sequencing is the price charged at the University of Florida (2020 pricing). The Next Generation Sequencing cost estimate is per student. The per student costs associated with the microscopes assumes that three students concurrently use each microscope in the below laboratories. Total costs for each laboratory are for the first time the laboratory is run. Costs would decrease after the initial running of the laboratories due to the presence of common equipment (i.e., the microscopes). The cost of common reagents required in the laboratories, facility upkeep/costs associated with obtaining laboratory space, service contracts for major equipment, and instruction delivery costs (Graduate Teaching Assistants), are not included in the below calculations. Importantly, animal costs are also not included in the below calculations.

| Equipment | Cost per Student |
| --- | --- |
| Dissecting microscope with camera | \$1,000 (\$3,000 microscope; prices vary widely based on specifications. Price is for base model) |
| Next generation sequencing sample preparation (3 replicates each for pooled fore- and hindlimbs) | \$600 |
| Next generation sequencing lane running (2 lanes per student; 4 samples/lane) | \$3,500 |
| Cost per student for major equipment/supplies | \$5,100 |

B. Laboratory 2: “The power of *C. elegans* in human gene discovery”,
| Equipment | Cost per Student |
| --- | --- |
| Florescent dissecting microscope | \$2,666 (\$8,000 microscope; prices vary widely based on specifications. Price is for base model) |
| Whole genome sequencing to identify mutations | \$500 |
| Cost per student for major equipment/supplies | \$3,166 |

C. Laboratory 3: “Investigation of regenerative capabilities”
| Equipment | Cost per Student |
| --- | --- |
| Dissecting microscope with camera | \$1,000 (\$3,000 microscope; prices vary widely based on specifications. Price is for base model) |
| Compound microscope to examine slides (with camera) | \$170 (\$510 microscope; prices vary widely based on specifications. Price is for base model) |
| Florescent dissecting microscope | \$2,666 (\$8,000 microscope; prices vary widely based on specifications. Price is for base model) |
| Cost per student for major equipment | \$3,836 |

To create mutations in *C. elegans*, animals are treated with ethyl methanesulfonate (EMS). EMS is a mutagenic and carcinogenic compound[21]. The use of the compound by undergrad students is prohibited so this lab could not be recreated in the residential program. In the virtual laboratory, *C. elegans* are incubated with EMS to induce mutations that potentially will alter the fate of M lineage cells (i.e. cells marked with GFP). The student picks male *C. elegans* and incubates them with EMS. Normally, to induce mutations in *C. elegans* DNA animals are treated for four hours, however, in an online environment this step only takes five seconds (Table 3b). The student then sets up a cross (males crossed to hermaphrodites) to generate offspring. While time is fast forwarded to allow progeny to grow to a stage where GFP expression can be scored for altered expression patterns, students are asked a series of questions to determine their understanding of the experiment they are performing. For example, the student is asked why the genetic cross they are performing does not result in all offspring being derived from the hermaphrodites (the correct answer is that male sperm outcompetes sperm present in hermaphrodites). Additional example questions are shown in Table 4.

**Table 3.** The total time each of the three laboratory experiments take to perform in the virtual laboratory simulations and if they have been performed in a brick-and-mortar environment. The time estimates are for performing experiments only. Analysis of results and experiment planning are not included.

| Experiment | Estimated time to perform on-ground | Estimated time to perform online |
| --- | --- | --- |
| Windowing/examination of stages of a chick egg | 7 days if egg lives<br>Experiment is halted and needs to be restarted if egg dies | 15 minutes<br>Experiment can be repeated as often is needed |
| Obtaining mouse embryos | 11 days from female becoming pregnant | Immediate<br>No mating, fertilization, or gestation required |
| <u>Total lab time needed</u> | 18 days | 15 minutes |

B. Laboratory 2: “The power of *C. elegans* in human gene discovery”
| Experiment | Estimated time to perform on-ground | Estimated time to perform online |
| --- | --- | --- |
| Identification of different stage *C. elegans* | 5 minutes | 5 minutes |
| Mutagenesis of *C. elegans* using EMS | 4 hours | 5 seconds |
| Genetic crosses to obtain recessive mutants | 5 days | 30 seconds |
| Expansion/purification of mutants | 3 weeks | Immediate |
| Screening for mutations | 2 days (very labor intensive; can take up to 15 hours over two days) | 1 minute |
| Whole genome sequencing to identify mutations | 7 days | Immediate |
| Confirmation of sequencing data* | 4 weeks | 30 seconds |
| <u>Total lab time needed</u> | <u>>2 months</u> | <u>~7 minutes</u> |

\* Confirmation of sequencing data includes sequencing specific regions of mutant and control strains and the performance of non-sequencing experiments to determine if a specific identified mutation is causing the observed mutant phenotype.

C. Laboratory 3: “Investigation of regenerative capabilities”
| Experiment | Estimated time to perform on-ground | Time to perform online |
| --- | --- | --- |
| Regeneration of an amputated axolotl limb | 40-50 days | 30 seconds |
| Regeneration of a sutured amputated axolotl limb | 40-50 days | 30 seconds |
| Regeneration of a GFP chimeric axolotl limb | 40-50 days | 30 seconds |
| <u>Total lab time needed</u> | <u>>4 months</u> | <u>~1 minute</u> |

**Table 4.** Sample questions and unedited feedback from students. **Questions** shown below were embedded in the online laboratories.

|  |  |  |
| --- | --- | --- |
| Sample Laboratory Questions | **Laboratory 1:** “Discovering genes responsible for patterning a vertebrate limb” | Which model organisms is best suited to visualize the development of forelimbs in a living vertebrate embryo? |
|  |  | What are the advantages of having the ability to watch the development of the same embryo? |
|  | **Laboratory 2:** “The power of *C. elegans* in human gene discovery” | Let's assume that the recessive mutation is viable and homozygous. What would you expect to see if we let a mutant F2 hermaphrodite self-fertilize? |
|  |  | How could we test that the identified gene really causes the observed phenotype? |
|  | **Laboratory 3:** “Investigation of regenerative capabilities” | Which of these blastemas is older? Explore, or take a look at them again. |
|  |  | You hypothesized before that the dedifferentiated cells are not able to multiply and that stem cells migrate from somewhere else. What pattern would you expect in the regenerating limb? |
| Sample Student Feedback | It was really interesting and new, it was a great way to see step by step processes that we'd experience in a lab. We talk a lot of processes but seeing them is different. |  |
|  | I really enjoyed the lab and wish we could have done more labs. It was a really hands-on experience without actually being hands on. I felt like I learned a lot more that way. I completely geeked out about windowing the chick eggs. It was super cool. |  |
|  | I loved the game-like interaction and that in turn, helps with learning and understanding certain concepts that otherwise might be missed in the lecture videos or assignments. For someone like me, a visual person, interaction helps me with the learning process. |  |
|  | It was great to work in a simulated development lab and to imagine what it would be like to think like a NGS scientist. In an in person lab it would be too expensive and time consuming to actually experience it. |  |
|  | The ability to "fast forward" and see what results would be like in a few moments that would take weeks to generate in an actual lab |  |
|  | It was fun to pretend I was doing the experiment! Especially since I'm not a fan of anything remotely lizard-like, the experiment allowed me to still experiment with a salamander whereas in an in person lab I would've run. |  |
|  | The best part of the lab, and with every lab we've done so far, is the ability to connect experiments done on model organisms (i.e, chick, nematode, and axolotl) and results with real world situations, especially in medicine. |  |
|  | I actually really enjoyed learning about all the mutant variations that scientists keep as a default for experiments. I feel like this really expanded my knowledge on real life applications better than any biology course I've taken so far (I'm a Junior). I really liked the labs. Usually I have a bit of anxiety in real life labs. Here, I could just learn and enjoy without worrying about consequences. |  |

This laboratory assumes the student has a basic understanding of genetics; however, genetics is not a prerequisite for enrolling in this course. To aid students in understanding the genetic crosses they need to perform, a real-time flow chart is provided in the online laboratory (Figure 2d). For example, animals generated in the first cross (referred to as the “F1” generation) would not be expected to have altered GFP expression unless a dominant mutation occurred. The student must perform a second cross to be able to observe recessive mutations. Animals from the second cross are examined under a florescent dissecting microscope to determine if there is a change in GFP expression. Each student analyzed two plates, with one plate containing animals with normal GFP expression and the other having altered GFP expression. The student is informed that their (virtual) laboratory assistant examined many other plates, with most of them containing normal M lineages.

To uncover the mutation responsible for the abnormal phenotype, the genome of the mutant animal is sequenced and the student is provided with a choice of three DNA changes that could be causing the abnormal phenotype. One of these DNA changes is in the coding region (i.e. changes the protein structure) of a FGFR gene, while the other two do not alter the proteins amino acid sequence. The laboratory concludes back in the hospital conference room where the team describes how the model system *C. elegans* was successfully used to identify new genes that may cause Saethre Chotzen Syndrome. When DNA from the case patient was sequenced, the child was found to have a mutation in the FGFR gene. In humans, Saethre Chotzen Syndrome patients that do not have a *twist* mutation often have mutations in *FGFR[22, 23]*.

In a residential setting, this laboratory would be extremely difficulty to accomplish since 1. EMS, a known carcinogenic is used, 2. the generation of EMS-induced mutations in an organism is random, 3. whole genome sequencing to identify mutations is relatively expensive, and 4. the time it would take to perform the experiments is immense. While it is possible that the instructor could generate EMS treated animals for each student, thus avoiding student exposure to EMS, manual screening of 1000’s of animals to identify the small number of animals containing a change in GFP expression would be too time consuming to accomplish during a typical laboratory period (Table 3b). One of the authors of this report performed a screen similar to the one described in the online laboratory[24]. In a pilot screen of ~1000 individual plates of *C. elegans*, one mutation affecting the M-linage was identified[24]. In addition, to score mutant animals, each student would require access to a florescent dissecting microscope resulting in the laboratory being prohibitively expensive to run (Table 2b). The identification of the mutation responsible for altered GFP expression in the EMS-induced animals would also be outside of most universities’ budgets[25–28] (Table 2b).

## Construction of the regeneration laboratory using the axolotl model system

In the regeneration laboratory, the learning objectives were:

1. Articulate why regeneration is important for survival of an organism.
2. Analyze regeneration in axolotls and interpret the results of transplantation experiments.
3. Describe why regeneration works poorly in humans.

The regeneration laboratory used the axolotl model to investigate the processes responsible for vertebrate regeneration. The laboratory began with a discussion of how some vertebrates can regenerate many structures, while other vertebrates (i.e. humans) have a limited capacity to regenerate tissues. The first experiment performed by the student recreates the classic axolotl limb amputation, which was reported in 1973[29, 30]. In this experiment, the student cuts off an axolotl limb and observes how it regenerates (the student cannot proceed without first anesthetizing the animal; Figure 3a). In real time, full regeneration of the limb takes ~50 days but in the online laboratory the student can watch complete regeneration of the limb in 35 seconds (Table 3c).

**Figure 3.**
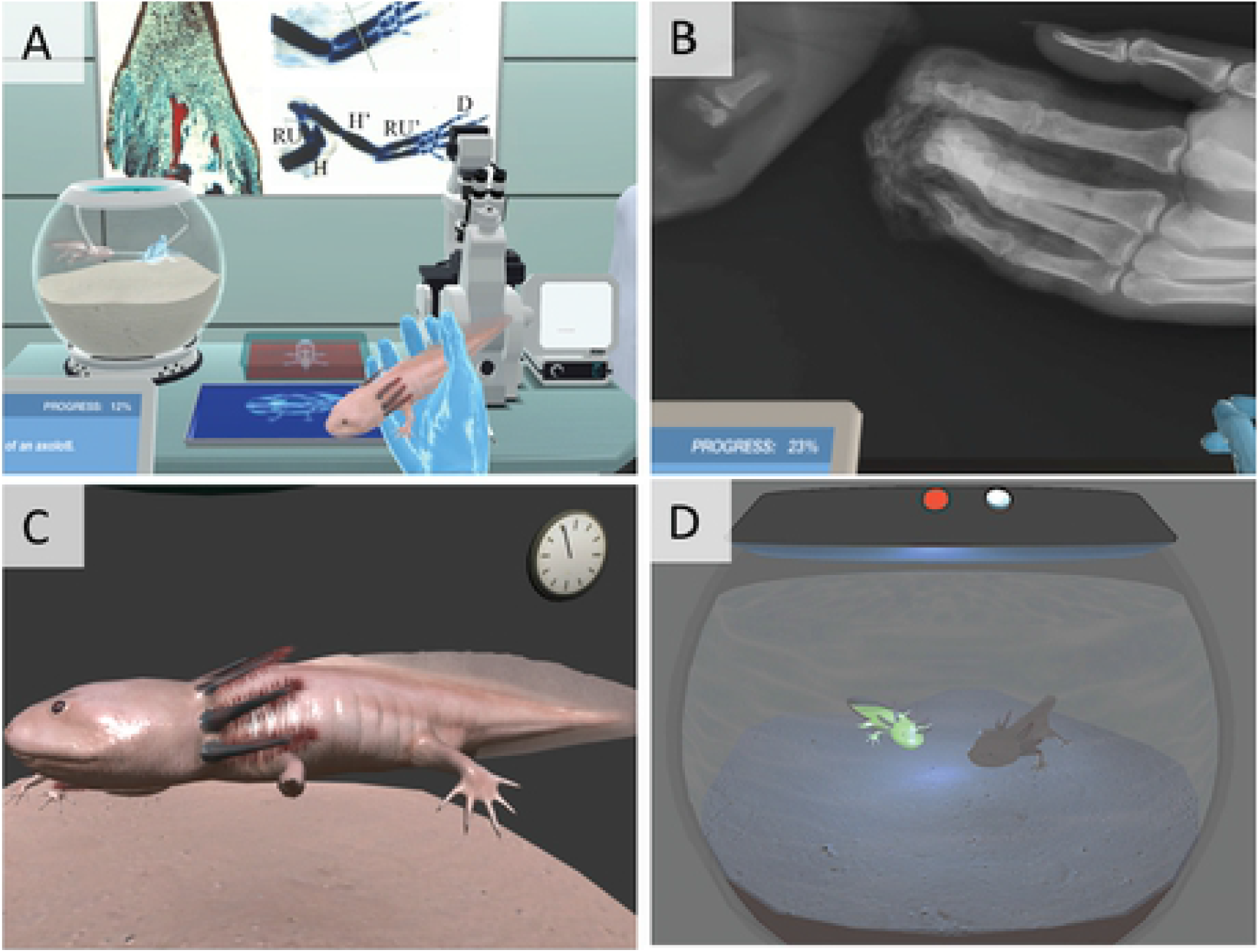
Images taken from Lab 3, “Investigation of regenerative capabilities”. (A) Dissection of an axolotl limb (the student cannot proceed without first anesthetizing the animal). (B) X-ray of a human limb containing amputation of three distal digits (C) Amputated followed by suturing of an axolotl limb. After ~50 days, or 35 seconds in the online laboratory, the student observes an absence of regeneration of the amputated sutured limb. (D) GFP-positive and GFP-negative axolotls.

An animated axolotl is used in the online laboratory. This was done to decrease the anxiety that some students may feel performing this experiment on “living” animals. As noted by one of the students in the course, they did not believe that they would have been able to complete the laboratory if they had been required to perform this experiment on a living animal (please see below and Table 4).

After completion of this initial experiment, the student is brought to the clinic where a young patient had just arrived at the hospital with amputated fingers. The student is shown an x-ray from this patient in which the patients bones were no longer connected to their hand (an x-ray from a human patient whose fingers were amputated is shown; Figure 3b). The student is then asked if they would like to see the corresponding picture of the amputation. Observation of a picture of the amputated fingers is not required to proceed. If the student decides that they would like to observe the picture, they are warned again about the graphic nature of the image. To observe the image of amputated fingers, the student must confirm twice that they understand that the image that are about to observe may be disturbing.

While two of the amputated human fingers were sutured, the third, which was amputated just above the nail bed, was not. The student is asked to propose a theory on why this single finger was left unrepaired. Any of the selected hypothesis could be correct and the student is promoted to return to the laboratory to determine why a physician would choose to leave an amputated finger untreated. The first hypothesis tested by the student is that suturing of a wound site inhibits regeneration. The student performs the axolotl amputation for a second time, with the single change that the wound is sutured after amputation. After ~50 days, or 35 seconds in the online laboratory, the student observes an absence of regeneration of the amputated sutured limb (Figure 3c). A morphological description, driven by the student correctly answering question, of the normal regeneration process in axolotls is then shown.

At this point in the laboratory, approximately two months have passed (the amount of “real time” that has passed is noted in the student’s notebook and they are promoted to check the number of days that have passed since they started their experiments; Table 3c). After 2 months, the initial patient who had amputated their fingers returns to the hospital for a checkup. A visualize examination of the patient’s hand reveals that the finger that was amputated distal to the nail bed, and that was not sutured, has regenerated while the fingers that had larger amputations did not regenerate. These observations lead to the second part of the laboratory where the student investigates what cells are responsible for regeneration.

In the second part of the laboratory, the student is shown tissue sections from two different stages of a regenerating axolotl limb. The student is lead through an analysis of the cell changes that occur during regeneration through the successful answering of a series of questions. These questions end with the hypothesis that the blastema (the part of the limb that is regenerating) can form all the necessary cell types to regenerate the limb. To test this hypothesis, the student is introduced to an axolotl in which all cells express GFP (Figure 3d). The student transplants a piece of skin from the GFP axolotl to a normal axolotl. This is a technically challenging experiment to perform in a residential laboratory. The chimeric animal contains a piece of GFP skin (i.e. these cell glow green when the protein is excited at the correct wavelength) while all other cells present in the animal lack GFP. The limbs of these animals are amputated and allowed to regenerate. Upon competition of regeneration, the student observes that not all cells present in the regenerated limb are GFP-positive (muscle cells are GFP-negative, while all other cells in the new limb are GFP-positive). These data indicate to the student that skin can form all cells of a regenerated limb except muscle cells.

The laboratory ends with the student playing a simulated “game”. In this simulation, the student is presented the skeleton of a limb and can amputate it in different regions. No matter where the amputation occurs, a normal new limb forms, indicating to the student that the limb has a mechanism in which it senses where the amputation occurs and only regenerates the part of the limb that is needed to form a normal structure. The second part of the simulation allows the student to perform the amputations in the presence of three molecules. This allows the student to begin to determine some of the molecular components that play a role in limb regeneration (amputation followed by regeneration in the presence of one of the molecules creates an abnormal longer limb, one has no effect, and one halts regeneration).

## Cost advantages of performing laboratories online

A major advantage of using virtual laboratories is that the cost of performing experiments can be ignored. Each of the three laboratories in this course would have been prohibitively expensive to perform in a face-to-face setting (Table 2). For example, the ability to house large numbers of mice and axolotls would severely tax the resources provided for face-to-face undergraduate laboratories. In both the Vertebrate Embryology Laboratory and Invertebrate Genetics Laboratory laboratories, each student performed large scale sequencing projects, which would not be possible in a brick-and-mortar setting.

The estimated cost to perform the Invertebrate Genetics Laboratory is >$3,166/student, the Vertebrate Embryology Laboratory >$5,100/student, and Vertebrate Regeneration Laboratory >$3,836/student (these costs are for the initial running of the laboratories). These costs assume that, if the laboratories were performed in a residential setting, each microscope is shared by three students. We note that costs could be modestly reduced by the sharing of common equipment across multiple laboratories (for example, a dissecting microscope is required in all three laboratories).

The costs noted above only take into consideration the major expenses needed to perform each laboratory. For example, the cost of basic laboratory supplies and expenses associated with maintenance of a physical laboratory space are not included in our calculations. In addition, the online course can enroll 50 students with a single instructor. The same number of students in the residential version of this course requires employment of an instructor and two teaching assistants.

## The ability to alter time online

A huge advantage of performing experiments online is the ability to alter time. Each of the laboratories would have been impossible to complete in a residential setting. For example, even the “shortest” lab, the Invertebrate Genetics Laboratory would have taken >18 days, of mostly full time laboratory work, to finish (our time estimate assumes an ideal situation where all experiments function correctly the first time; Table 3). Online, each laboratory was designed to be accomplished in <60 minutes.

In online laboratories, students have the ability to not only speed up time but to slow it down. In our online laboratories, we did not take advantage of the capability to perform an experiment in slow motion, but this ability would be ideal for experiments that proceed rapidly in real-time (for example, some chemical reactions). In addition, online laboratories allow students the ability to go back in time to re-perform an experiment or check their results.

## Assessment of laboratory learning outcomes

All three virtual laboratories in the course were anchored around a human disease/problem. In addition, the Vertebrate Embryology Laboratory began in a hospital setting since many of the students in the course were interested in medically-related careers. Feedback from the students (Table 4) suggested that the students found the laboratory content engaging and medically relevant.

Each laboratory was designed to be performed in approximately 60 minutes. Students were allowed to submit the completed virtual laboratory once for a grade, but were able to start, stop, and restart the lab session as many times as they wished without penalty (up until the due date). This was purposefully designed so students could make mistakes, pause and consult their lecture notes, each other, or their instructor if they found themselves feeling unprepared or confused at any point in the virtual laboratories. The final score obtained in the virtual laboratories pushed to the learning management system’s gradebook tool once a student submitted the final question in the laboratory.

Throughout each laboratory, questions were strategically embedded to test comprehension (15-26 questions per laboratory). Students were encouraged to access background theory present in the virtual laboratories and to use what they learned as they answered each question. Students could attempt each 10-point question up to four times, losing two points per attempt. Between each attempt, answer choices were reordered and real-time feedback was provided (through the simulation). Once the virtual laboratory was submitted, students were given a quiz that tested their understanding of the laboratory (outside of the online laboratory simulation). The quiz required the student to use the information they learned in the virtual laboratory to successfully answer the quiz questions.

## Conclusion and the future of online science laboratories

The laboratories described in this report would not have been possible to perform in a residential setting. These laboratories would have been too expensive, time consuming, and in some cases dangerous for students to perform (Tables 1-3). In addition, the creation of mutations in forward genetic screens is random making it extremely difficult to ensure a relatively uniform laboratory experience for each student. Online, none of these variables are an issue.

Five years-ago the creation of the laboratories described in this report would have likely not been possible due to technological roadblocks and more importantly a lack of expertise (and interest) in creating virtual undergraduate laboratories. Due to the creation of 100% online degrees, the University of Florida has taken an innovative and exciting approach to the delivery of online laboratories. It is important to note that the University of Florida, and we believe most centers for higher education, lack in-house resources to create 100% online laboratories that can fulfill the requirements of a science degree. A collaboration between Labster, a company based in Denmark that specializes in simulating an online laboratory setting and the authors of this report produced an environment in which online students had the opportunity meet the same learning outcomes as residential students.

A common complaint about the online laboratory experience is that students are unable to learn and practice many of the physical techniques commonly used in a laboratory setting. The authors of this report have extensive experience mentoring undergraduate and graduate students in their laboratories over decades and believe that it is the ability to generate hypothesis, develop a way to test them, and analyze the resulting outcomes that are the essential skills laboratory courses should be designed to teach. In our experience, the ability to master psychomotor learning objectives like slide preparation techniques is not the most important aspect of a laboratory experience. Our experience designing and teaching the online laboratories described in this report suggests that upper division online laboratories are not only feasible but have the potential to exceed the learning outcomes of their residential counterparts.

The development of 100% online courses (and degrees) at UF pre-date the COVID19 pandemic. As noted previously, online laboratory experiences at UF were developed to provided 100% online undergraduate degrees to students in UF Online. In the case of the online laboratories described in this report, these laboratories had already been incorporated into the residential version of the course due to the positive experiences reported by the online students. As the entire world transitioned to a mostly online modality in Spring 2020, UF used previously developed 100% online courses, including the one described in this report, to allow residential students to continue to make progress towards their degree.

## Acknowledgements

The authors would like to thank UF Online for providing funding to create the three virtual online laboratories. Strong support from UF Online administration was essential for the successful development of these laboratories. The conversion of our ideas into a virtual simulation would not have been possible without the abilities of Labster. Andrew McDonald (UF Information Technology) played a key role in integrating the finished laboratories into the course. The current paper would not have been possible without the willingness of UF undergraduates to try something new.

## References

1. Hallyburton, C.L. and E. Lunsford, Challenges and Opportunities for Learning Biology in Distance-Based Settings. Bioscene: Journal of College Biology Teaching, 2013. v39(1): p. p27–33.

2. Rutten, N., R.W. van Joolingen, and J. van der Veen, The learning effects of computer simulations in science education. Computers and Education, 2012. 58(1): p. 136–153.

3. Senate, T.F., Appropriations subcommittee on Education (C/CS/SB1076:K-20 Education), Tallahassee, FL.

4. Tiberio, G., et al., Liebenberg syndrome: brachydactyly with joint dysplasia (MIM 186550): a second family. J Med Genet, 2000. 37(7): p. 548–51.

5. Seoighe, D.M., et al., A chromosomal 5q31.1 gain involving PITX1 causes Liebenberg syndrome. Am J Med Genet A, 2014. 164A(11): p. 2958–60.

6. Spielmann, M., et al., Homeotic arm-to-leg transformation associated with genomic rearrangements at the PITX1 locus. Am J Hum Genet, 2012. 91(4): p. 629–35.

7. Al-Qattan, M.M., et al., Liebenberg syndrome is caused by a deletion upstream to the PITX1 gene resulting in transformation of the upper limbs to reflect lower limb characteristics. Gene, 2013. 524(1): p. 65–71.

8. Cherner, Y., et al., Use of adaptive simulation-based virtual laboratories for teaching alternative energy and energy conservation in engineering and technology programs. American Society for Engineering Education. 14(26): p. 10481–10492.

9. Rock, J.R., et al., Identification of genes expressed in the mouse limb using a novel ZPA microarray approach. Gene expression patterns: GEP, 2007. 8(1): p. 19–26.

10. Logan, M. and C.J. Tabin, Role of Pitx1 upstream of Tbx4 in specification of hindlimb identity. Science, 1999. 283(5408): p. 1736–9.

11. Naiche, L.A. and V.E. Papaioannou, Loss of Tbx4 blocks hindlimb development and affects vascularization and fusion of the allantois. Development, 2003. 130(12): p. 2681–93.

12. Duboc, V. and M.P. Logan, Pitx1 is necessary for normal initiation of hindlimb outgrowth through regulation of Tbx4 expression and shapes hindlimb morphologies via targeted growth control. Development, 2011. 138(24): p. 5301–9.

13. Mennen, U., S. Mundlos, and M. Spielmann, The Liebenberg syndrome: in depth analysis of the original family. J Hand Surg Eur Vol, 2014. 39(9): p. 919–25.

14. Gallagher, E.R., C. Ratisoontorn, and M.L. Cunningham, Saethre-Chotzen Syndrome, in GeneReviews(R), R.A. Pagon, et al., Editors. 1993: Seattle (WA).

15. el Ghouzzi, V., et al., Mutations of the TWIST gene in the Saethre-Chotzen syndrome. Nat Genet, 1997. 15(1): p. 42–6.

16. Howard, T.D., et al., Mutations in TWIST, a basic helix-loop-helix transcription factor, in Saethre-Chotzen syndrome. Nat Genet, 1997. 15(1): p. 36–41.

17. Brenner, S., The genetics of behaviour. Br Med Bull, 1973. 29(3): p. 269–71.

18. Ankeny, R.A., The natural history of Caenorhabditis elegans research. Nat Rev Genet, 2001. 2(6): p. 474–9.

19. Chalfie, M., et al., Green fluorescent protein as a marker for gene expression. Science, 1994. 263(5148): p. 802–5.

20. Harfe, B.D., et al., Analysis of a Caenorhabditis elegans Twist homolog identifies conserved and divergent aspects of mesodermal patterning. Genes Dev, 1998. 12(16): p. 2623–35.

21. Sega, G.A., A review of the genetic effects of ethyl methanesulfonate. Mutat Res, 1984. 134(2-3): p. 113–42.

22. Paznekas, W.A., et al., Genetic heterogeneity of Saethre-Chotzen syndrome, due to TWIST and FGFR mutations. Am J Hum Genet, 1998. 62(6): p. 1370–80.

23. Freitas, E.C., et al., Q289P mutation in FGFR2 gene causes Saethre-Chotzen syndrome: some considerations about familial heterogeneity. Cleft Palate Craniofac J, 2006. 43(2): p. 142–7.

24. Harfe, B.D., et al., MyoD and the specification of muscle and non-muscle fates during postembryonic development of the C. elegans mesoderm. Development, 1998. 125(13): p. 2479–88.

25. Zuryn, S. and S. Jarriault, Deep sequencing strategies for mapping and identifying mutations from genetic screens. Worm, 2013. 2(3): p. e25081.

26. Sarin, S., et al., Caenorhabditis elegans mutant allele identification by whole-genome sequencing. Nat Methods, 2008. 5(10): p. 865–7.

27. O’Rourke, S.M., et al., Rapid mapping and identification of mutations in Caenorhabditis elegans by restriction site-associated DNA mapping and genomic interval pull-down sequencing. Genetics, 2011. 189(3): p. 767–78.

28. Smith, H.E., et al., Mapping Challenging Mutations by Whole-Genome Sequencing. G3 (Bethesda), 2016. 6(5): p. 1297–304.

29. Stocum, D.L., Stages of forelimb regeneration in Ambystoma maculatum. J Exp Zool, 1979. 209(3): p. 395–416.

30. Iten, L.E. and S.V. Bryant, Forelimb regeneration from different levels of amputation in the newt,Notophthalmus viridescens: Length, rate, and stages. Wilhelm Roux Arch Entwickl Mech Org, 1973. 173(4): p. 263–282.

